# Hepatotoxicity during 6-thioguanine treatment in inflammatory bowel disease and childhood acute lymphoblastic leukaemia: a systematic review

**DOI:** 10.1101/535518

**Authors:** Linea Natalie Toksvang, Magnus Strøh Schmidt, Sofie Arup, Rikke Hebo Larsen, Thomas Leth Frandsen, Kjeld Schmiegelow, Cecilie Utke Rank

## Abstract

**Background:** The recently established association between higher levels of DNA-incorporated thioguanine nucleotides and lower relapse risk in childhood acute lymphoblastic leukaemia (ALL) calls for reassessment of prolonged 6-thioguanine (6TG) treatment, while avoiding the risk of hepatotoxicity.

**Objectives:** To assess the incidence of hepatotoxicity in patients treated with 6TG, and to explore if a safe dose of continuous 6TG can be established.

**Data sources:** Databases, conference proceedings, and reference lists of included studies were systematically searched for 6TG and synonyms from 1998–2018.

**Methods:** We included studies of patients with ALL or inflammatory bowel disorder (IBD) treated with 6TG, excluding studies with 6TG as part of an intensive chemotherapy regimen. We uploaded a protocol to PROSPERO (registration number CRD42018089424). Database and manual searches yielded 1823 unique records. Of these, 395 full-texts were screened for eligibility. Finally, 134 reports representing 42 studies were included.

**Results and conclusions:** We included data from 42 studies of ALL and IBD patients; four randomised controlled trials (RCTs) including 3,993 patients, 20 observational studies including 796 patients, and 18 case reports including 60 patients. Hepatotoxicity in the form of sinusoidal obstruction syndrome (SOS) occurred in 9–25% of the ALL patients in two of the four included RCTs using 6TG doses of 40–60 mg/m^2^/day, and long-term hepatotoxicity in the form of nodular regenerative hyperplasia (NRH) was reported in 2.5%. In IBD patients treated with 6TG doses of approximately 23 mg/m^2^/day, NRH occurred in 14% of patients; SOS has not been reported. At a 6TG dose of approximately 12 mg/m^2^/day, NRH was reported in 6% of IBD patients, which is similar to the background incidence. According to this review, doses at or below 12 mg/m^2^/day are rarely associated with notable hepatotoxicity and can probably be considered safe.

## INTRODUCTION

Hepatotoxicity was first noted as a complication of 6-thioguanine (6TG) treatment in the late 1970s[1] and has since been described in the form of sinusoidal obstruction syndrome (SOS) in childhood acute lymphoblastic leukaemia (ALL)[2–4] and nodular regenerative hyperplasia (NRH) in both inflammatory bowel disease (IBD)[5] and childhood ALL.[6] This reflects that SOS and NRH possibly represent a spectrum of related hepatic microvascular disorders spanning acute onset to chronic manifestation, although the underlying cellular mechanisms that connect them remain to be established Fig 1.[7–9]

**Fig 1.**
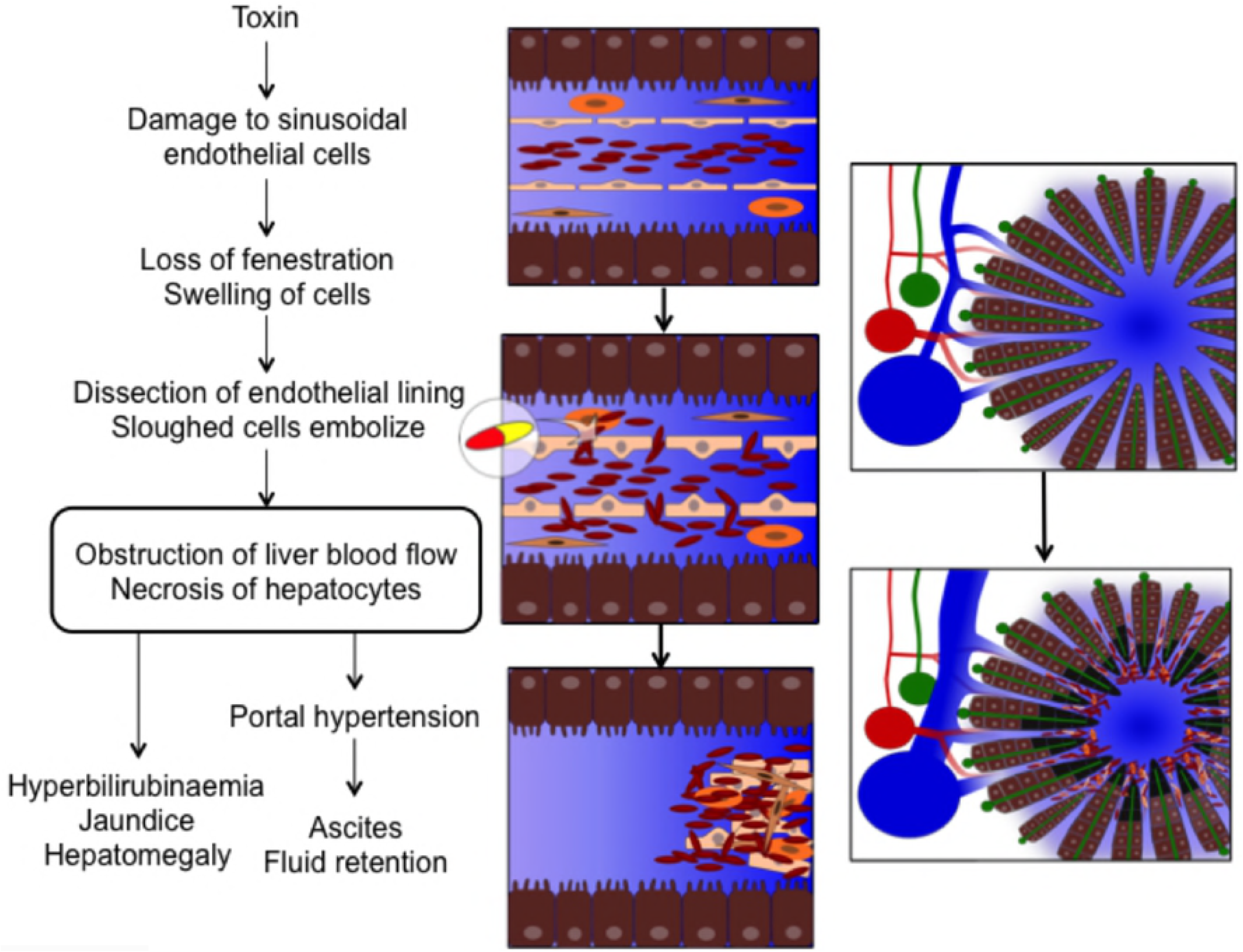
The pathophysiology of sinusoidal obstruction syndrome. The presumed pathophysiological mechanism underlying sinusoidal obstruction syndrome is a drug-mediated damage to sinusoidal endothelial cells, causing swelling and loss of fenestration. This allows red blood cells to enter the space of Disse and dissect off the endothelial lining. The sloughed off cells embolise downstream and cause obstruction of the hepatic microcirculation and consequently hepatocellular necrosis.[7] These pathophysiological changes lead to the clinical symptoms: jaundice, tender hepatomegaly, ascites and fluid retention. Figure by Linea Natalie Toksvang.

The development and severity of 6TG-induced hepatotoxicity appear to be dose-related, and rarely occurs at low cumulative doses.[10,11] Hence, a non-systematic review from 2013 focusing on IBD patients reported NRH in 0–53% of patients at 6TG doses of 12–58 mg/m^2^/day (20–100 mg/day) with no cases reported with 6TG doses below 12 mg/m^2^/day.[5] A meta-analysis from 2011 analysed 6TG treatment in childhood ALL and reported a seven-fold increased incidence of SOS in two of three randomised controlled trials (RCTs), in which patients received 6TG at doses of 40–60 mg/m^2^/day instead of 6-mercaptopurine (6MP) during maintenance treatment.[12]

Overall survival for childhood ALL now exceeds 90%;[13] however the poor prognosis following relapse still remains a major challenge.[14] Thus, although a number of novel anti-leukaemic treatment options, including immunotherapy, have emerged in recent years, their role in first line therapy remains uncertain.[13] Therefore, it is important to explore possible improvements with the use of the traditional anti-leukaemic agents, including thiopurines. The theoretical pharmacological advantages of 6TG over 6MP include a more direct intracellular activation pathway, higher potency, and shorter duration of exposure necessary for cytotoxicity. The cytotoxic effect of 6TG and 6MP is ascribed to the metabolite thioguanine nucleotides (TGN), which are incorporated into DNA and RNA as false purine bases as well as inhibiting de novo synthesis of purines.[3,15] Red blood cell (RBC) levels of TGN (ery-TGN) have been assumed to reflect treatment intensity and adherence. An ery-TGN level of more than 230–260 pmol/8·10^8^ RBC has been associated with therapeutical response in IBD patients. However, the association between metabolite levels, efficacy, and adverse events remains uncertain.[16] It was recently demonstrated that ery-TGN correlate with TGN incorporated into leucocyte DNA (DNA-TGN). Furthermore, higher levels of DNA-TGN correlate with a lower relapse risk in childhood ALL.[17] This finding calls for a reassessment of the feasibility of prolonged 6TG treatment for childhood ALL, while avoiding the risk of severe hepatotoxicity.

The primary objective of this systematic review was to assess the incidence of hepatotoxicity in IBD and childhood ALL patients treated with 6TG compared to 6MP or standard of care. The secondary objective was to explore if a safe dose of 6TG can be established.

## METHODS

We conducted this systematic review in accordance with the Cochrane Handbook[18] and Preferred Reporting Items for Systematic Reviews and Meta-Analyses (PRISMA) statement.[19,20] The systematic review protocol was uploaded to the International Prospective Register of Systematic Reviews (PROSPERO) before the literature search was conducted (registration number CRD42018089424), and is presented in S1 Appendix. The review authors were not blinded to the journal titles, study authors or institutions. In case of insufficient reporting, study authors were contacted for additional information. We sought to avoid double counting by juxtaposing author names, location, setting, and sample sizes.

### Eligibility criteria

We included RCTs, controlled clinical trials including quasi-randomised trials, observational studies (e.g. prospective and retrospective cohort studies, case-control, and cross-sectional studies), case series, and case reports in which patients at any age were treated with 6TG. We excluded studies investigating 6TG as part of an intensive chemotherapy regimen. Since most studies using 6TG for an extended period of time either as monotherapy or as part of maintenance treatment are within the field of IBD and ALL, we chose to limit the review to these two disease populations. We compared 6TG treatment to 6MP or standard of care (other non-6TG treatment regimens). We included studies published between 1998 and March 2018. We made no restrictions regarding timing, type of setting or language.

### Information sources

We systematically searched PubMed/MEDLINE, Embase/Ovid, Scopus/Elsevier, Web of Science Core Collection/Clarivate Analytics, and The Cochrane Central Register of Controlled Trials (CENTRAL)/Wiley (The Cochrane Library, issue 2, 2018). The search strategy included synonyms of 6TG in title, author keywords or thesaurus, and the time frame 1998 to 2018, and was conferred with a health care librarian. The specific search strategies for each database are presented in S2 Appendix.

Additional manual searches included reference lists of included studies, conference proceedings from the American Society of Hematology (ASH), European Hematology Association (EHA), American College of Gastroenterology (ACG), European Crohn’s and Colitis Organization (ECCO), American Association for the Study of Liver Diseases (AASLD), International Union of Basic & Clinical Pharmacology (IUPHAR), and European Association for Clinical Pharmacology and Therapeutics (EACPT). The ClinicalTrials.gov registry (www.clinicaltrials.gov), the International Standard Randomized Controlled Trial Number (ISRCTN) registry (www.controlled-trials.com), and the WHO International Clinical Trials Registry Platform (ICTRP) (www.who.int/ictrp/search/en) were searched for research protocols, and PROSPERO was searched for on-going systematic reviews. We conducted the latest search on March 7, 2018.

### Study selection

We used Covidence systematic review software (Veritas Health Innovation, Melbourne, Australia) to remove duplicates and conduct the screening. Two review authors independently screened titles and abstracts and subsequently evaluated eligible reports in full text. We recorded reasons for exclusion.

### Data collection process

Two review authors independently extracted data from included studies in duplicate using a standardised extraction form, which was approved by all authors and pilot tested on ten reports. A third author checked data extraction. A simplified sheet was used for articles in Polish, Russian, and Chinese – with consultant help for data extraction. Extracted data items included authors and year of publication, study characteristics (design, blinding, setting, duration, inclusion and exclusion criteria, source of funding, conflict of interest, ethical approvals, key conclusions), patient characteristics (number of patients, age, sex, ethnicity, disease, comorbidity, concomitant therapy), details of the interventions (duration of 6TG, dose of 6TG, cumulative dose of 6TG, maximum dose of 6TG, route of administration, ery-TGN levels), comparator (comparator drug, duration of 6MP or other standard of care drug, dose of 6MP or other standard of care drug, cumulative dose of 6MP or other standard of care drug, maximum dose of 6MP or other drug, route of administration), follow-up, and outcomes as described below.

#### Primary outcome

Incidence of any hepatotoxicity reported as SOS, veno-occlusive disease (VOD), NRH, discordant thrombocytopenia (transfusion-resistant and/or unexplained by treatment), drug-induced liver injury or non-specified hepatotoxicity. Due to the lack of standardised definitions of hepatotoxicity, authors of the included studies may not have used the above-mentioned terms. To assess additional hepatotoxicity we therefore report any pathological findings of liver biopsies and use the *Ponte di Legno* (PdL) toxicity working group consensus criteria for SOS, which entail fulfilment of at least three out of the following five criteria: (i) hyperbilirubinaemia; (ii) hepatomegaly; (iii) ascites; (iv) weight gain of 5% or more; (v) discordant thrombocytopenia.[21] Furthermore, we considered an increase in alanine transaminase, aspartate transaminase, alkaline phosphatase, conjugated bilirubin or total bilirubin of more than two times upper normal limit as evidence of hepatotoxicity.[22]

#### Secondary outcomes

Diagnostic methods (number of patients who had a liver biopsy, indication for liver biopsy, other diagnostic methods, study conclusions about diagnostic methods); dose reduction or truncation of 6TG due to hepatotoxicity (how many patients had 6TG truncated or the dose reduced; doses before and after dose reduction).

### Risk of bias of individual studies

Two review authors independently evaluated risk of bias of the included reports using the Cochrane Collaboration tool for RCTs. [18] The study quality assessment tools of the National Heart, Lung, and Blood Institute of the National Institutes of Health (NIH) for quality assessment of Observational Cohort and Cross-Sectional Studies and Controlled Intervention Studies[23] were used to assess the risk of bias of observational studies. A graphic illustration of potential bias of RCTs was made using the Review Manager (RevMan [Computer program]. Version 5.3. Copenhagen: The Nordic Cochrane Centre, The Cochrane Collaboration, 2014). Studies with low risk of bias were ascribed more weight in the overall findings of the review compared to studies with higher risk of bias.

### Assessment of heterogeneity

We will visualise the findings of the included studies and heterogeneity between the studies in a figure. Meta-regression analyses were not performed.

### Data synthesis

We present a systematic narrative synthesis of the characteristics and findings of the included studies, since the studies were too heterogeneous in terms of design and comparator in order to perform a quantitative data synthesis. We calculated doses in mg/m^2^ with the assumption that an average adult is 1.73 m^2^, and that 30 kg correspond to one m^2^. Data are presented as median (range) unless otherwise stated.

### Meta-biases

We addressed publication bias by searching for and including grey literature, such as conference abstracts not published in journals or books. To address selective outcome bias, we compared outcomes between protocols and published reports. If protocols were unavailable, we compared outcomes reported in the methods and results sections. We did not quantitate the impact of meta-biases.

### Confidence in cumulative evidence

The Grading of Recommendations, Assessment, Development and Evaluation (GRADE) approach[24] was used to assess the confidence in cumulative evidence. Two review authors independently evaluated the quality of evidence in each of four domains: (1) study design, (2) study quality, (3) consistency, and (4) directness. The evidence on each outcome was graded as ‘very low’, ‘low’, ‘moderate’ or ‘high’.

## RESULTS

### Study selection

A flow diagram of study selection is presented in Fig 2. The database and manual searches yielded 1823 unique records. Of these, we screened 395 in full-text for eligibility. Finally, 134 reports representing 42 studies were included in the review. Of these, four were RCTs [2,3,25,26], 20 were observational studies (6,27–45), and 18 were case reports (11,46–62). No similar reviews were found in the database search or on PROSPERO.

**Fig 2.**
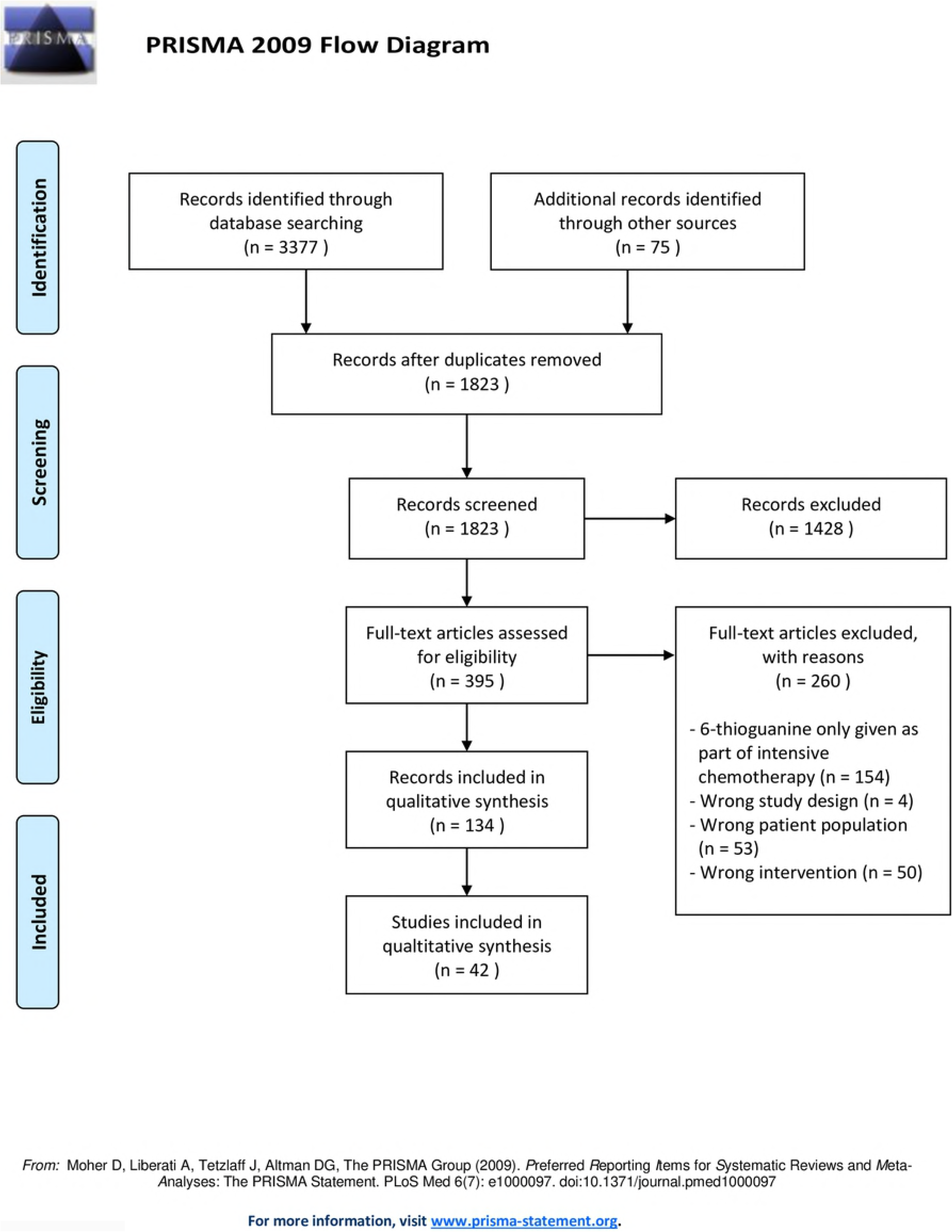
PRISMA Flow diagram of study selection.

### Study characteristics

Characteristics of the individual studies are presented in S3 Appendix.

### Risk of bias within studies

Judgements of each risk of bias item for included RCTs are illustrated in S4 Appendix. None of the RCTs were blinded in respect to participants and personnel or outcome assessment. Risk of bias evaluations of observational studies are included in Table B in S3 Appendix. No serious bias was suspected, yet in most studies outcome assessors were not blinded to exposure status, and no studies used power calculations to justify sample size.

### Meta-biases

Manual searches yielded additionally two reports eligible for inclusion in the review. Six conference abstracts without a published original article were found, but the authors did not respond to our enquiries. Risk of bias was not evaluated for conference abstracts without an associated article. Protocols were only available for two RCTs[2,3] and one pharmacokinetic study,[34] in which no inconsistencies were found. Outcomes reported in the methods and results sections of the remaining reports were compared, and no serious inconsistencies were found.

### Confidence in cumulative evidence

The confidence in cumulative evidence evaluation is presented in Table 1.

**Table 1.** Overview of GRADE evaluation.

| Outcomes | Study design, n | Study quality | Consistency | Directness | Overall quality |
| --- | --- | --- | --- | --- | --- |
| **Incidence of hepatotoxicity** | Randomised clinical trials, 4<br>Observational studies, 15<br>Case reports, 14 | No serious limitations | Important inconsistency in the definition of hepatotoxicity and how it is described | No uncertainty about directness | Moderate |
| **Diagnostic methods** | Randomised clinical trials, 2<br>Observational studies, 8<br>Case reports, 5 | No serious limitations | Important inconsistency in the use of diagnostic modalities across studies | No uncertainty about directness | Moderate |
| **Reduction or truncation of 6TG** | Randomised clinical trials, 3<br>Observational studies, 10<br>Case reports, 9 | No serious limitations | No serious inconsistency | No uncertainty about directness | High |

## Results of individual studies

Fig 3 represents the incidence of SOS or NRH compared to 6TG dose in the included studies. A detailed overview of the results of the individual studies is presented in S3 Appendix.

**Fig 3.**
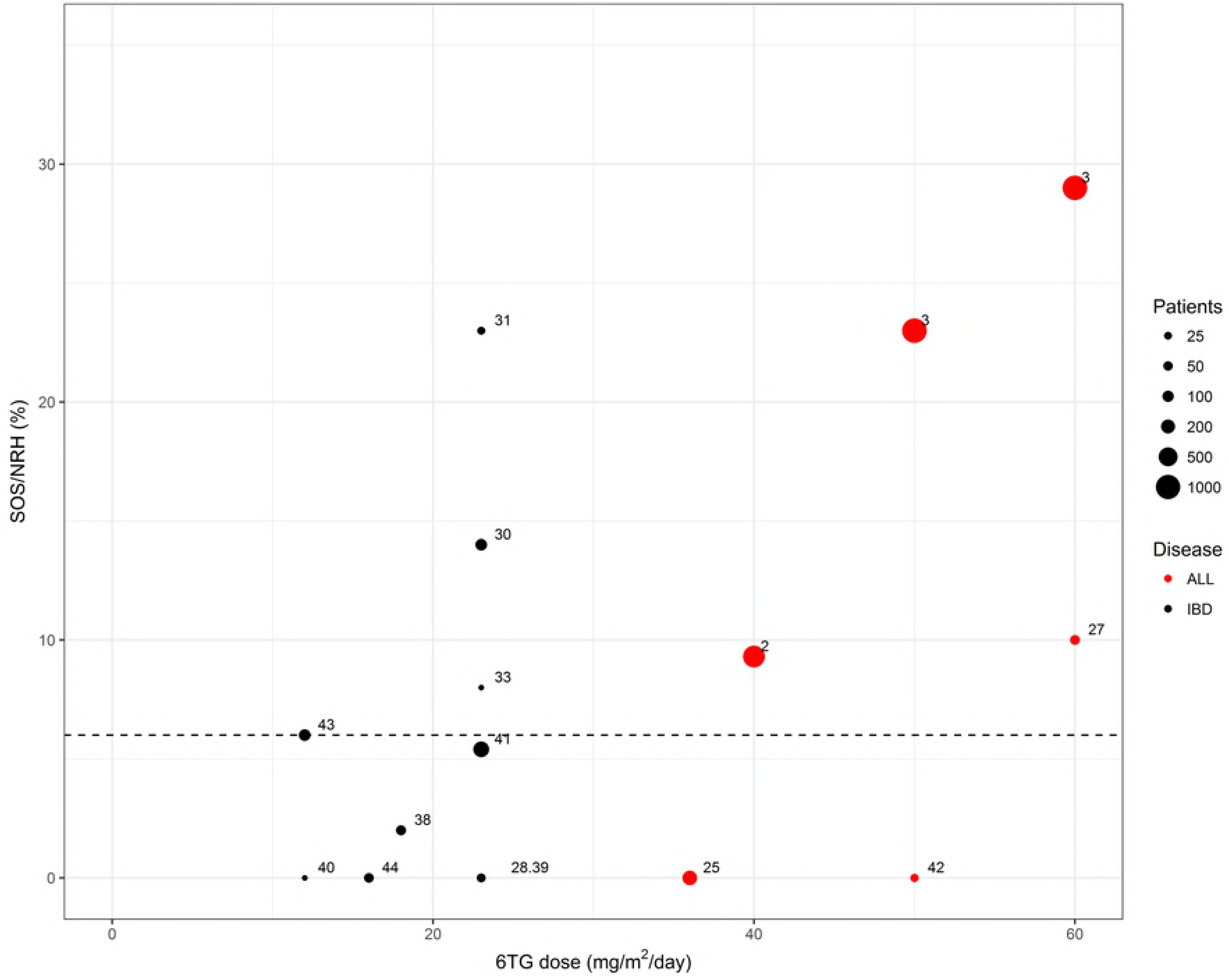
Incidence of sinusoidal obstruction syndrome (SOS) or nodular regenerative hyperplasia (NRH) compared to 6-thioguanine (6TG) dose in the included studies. Doses in mg/m^2^ were calculated with the assumption that an adult is 1.73 m^2^, and that 30 kg correspond to one m^2^.

### Synthesis of results

We included data from a total of 42 studies, encompassing four RCTs including 3,993 patients, 20 observational studies including 796 patients, and 18 case reports including 60 patients.

In two of three RCTs on 6TG versus 6MP at doses 6TG of 40–60 mg/m^2^/day for maintenance treatment in childhood ALL, hepatotoxicity was reported in the form of SOS in up to 25% of patients in the 6TG-arm.[2,3] The incidence of SOS was highest in the CCG-1952 trial, which used the highest target dose. Moreover, in this trial the incidence of SOS was reduced to 20% when the 6TG target dose was changed from 60 to 50 mg/m^2^.[3] Evidence of persisting hepatotoxicity including NRH and complications of portal hypertension were reported in 2.5% of patients receiving 6TG. No hepatotoxicity was reported during 6MP treatment.[2,3] One of the RCTs on 6TG versus 6MP did not report SOS, but found evidence of hepatotoxicity by means of an increased risk of discordant thrombocytopenia during 6TG treatment.[25]

Starting doses of 6TG of approximately 23 mg/m^2^/day were used in five observational studies of adults with IBD, which were associated with elevated liver enzymes in 8–25% of patients and histological evidence of NRH in 14–23% of patients. Seven studies used a lower 6TG dose of approximately 12 mg/m^2^/day, and NRH was reported in 0–6% of patients.[43]

In regard to the included case reports, the 6TG dosage was higher in the ones that reported hepatotoxicity (14–125 mg/m^2^/day)[47,48,59,62,49–53,55,57,58] compared to the ones that did not report hepatotoxicity (0.8–21 mg/m^2^/day).[41,54,56,61]

Hepatotoxicity documented by blood samples, various imaging modalities, or liver biopsy was not persistently associated with higher ery-TGN levels.[30,63,64]

The included RCTs were generally well executed with low risk of bias, except in the areas of blinding participants and outcome assessment, which may lead to performance and detection bias. Risk of bias assessments in the included observational studies did not result in any suspicion of specific biases. The confidence in the cumulative estimate of the incidence of hepatotoxicity was graded as moderate, and the use of diagnostic methods as moderate, primarily because of the inconsistencies in reporting within these subjects. The confidence in the cumulative estimate of dose reduction or truncation of 6TG was graded as high.

## DISCUSSION

The findings of the present systematic review indicate that 6TG-induced hepatotoxicity in the form of SOS or NRH is highly dose-dependent, and that it rarely occurs at daily doses of less than 12 mg/m^2^/day. Furthermore, 6TG-induced hepatotoxicity appears to be largely reversible, except at high doses exceeding 40 mg/m^2^/day.

The three RCTs of childhood ALL, the COALL-92 trial, the CCG-1952 trial, and the UK MRC ALL97/99 trial, respectively, have previously been included in a meta-analysis, which estimated the increase in SOS between treatment arms to be a factor 7.16 (OR, 95% CI 5.66–9.06).[12] SOS was highlighted as a dose-related toxicity with a multifactorial aetiology. No SOS was reported in the COALL-92 trial, albeit a high frequency of discordant thrombocytopenia, which may reflect a minor degree of hepatotoxicity. In that trial, vincristine and corticosteroids were not co-administered with 6TG – as was the case in the other two RCTs.[25] This difference in the co-medication has been proposed to be an explanation for the differing incidence of SOS between the three RCTs.[2]

The theory that NRH is 6TG dose or ery-TGN level-dependent was first presented in 2005.[11] This is supported by a mouse model, in which SOS arose from high peak concentrations of 6TG, but did not occur with low daily doses of 6TG.[10] Most of the included studies did not find an association between ery-TGN-levels and hepatotoxicity, thus the findings do not support the use of ery-TGN for assessing the risk of 6TG-induced hepatotoxicity.

Several factors in the included observational studies investigating IBD may have led to an overestimation of hepatotoxicity. The risk of NRH development is associated with the use of immunosuppressive drugs often applied in IBD such as azathioprine, 6MP, ciclosporin, and steroids.[65] Thus, hepatotoxicity has been recognised in as many as 20% of IBD patients treated with azathioprine, 6MP, or methotrexate.[66] Azathioprine is the thiopurine most often associated with NRH,[67] and since most of the 6TG-treated IBD patients have received azathioprine previously, signs of NRH could have been present beforehand. As baseline liver biopsies prior to 6TG treatment were not obtained in any of the studies, some of the reported cases of NRH may likely have been triggered by other factors than 6TG. Furthermore, NRH or sinusoidal dilatation has previously been found in 6% and 34% of liver biopsies, respectively, from thiopurine-naïve IBD patients,[68] and the prevalence of NRH is probably as high as 5% in the background population, as suggested by autopsy studies of otherwise healthy individuals.[65,69] Thus, the incidence of NRH around 6% in the studies using 6TG at doses of approximately 12 mg/m^2^/day may not differ notably from the background incidence. Additionally, selection bias arising from only taking a biopsy in patients with suspected hepatotoxicity is supposedly present in most of the studies included in this review. Only one study performed liver biopsies in all patients in the cohort, avoiding selection bias, and found NRH in 6 % (N=7).[43]

The patient populations of the observational IBD studies had all failed conventional immunosuppressive treatment because of clinical unresponsiveness or adverse events, and undergone substantial surgery. Thus, the results cannot be generalised to the broader group of IBD patients. Moreover, 6TG undergoes extensive first-pass metabolism in the intestines and the liver; yielding bioavailability between 5–37%.[29] How IBD per se affects absorption and pharmacokinetics of 6TG is unknown.[34] The true incidence of hepatotoxicity in IBD patients treated with 6TG therefore remains unknown. However, a consensus paper by the European 6TG working party from 2006 recommends that 6TG can be used in a clinical research setting, in doses not exceeding 25 mg/day, corresponding to 14 mg/m^2^/day, which is considered a safe dose.[70] This is supported by the results of this systematic review.

While liver biopsy with reticulin stain remains the gold standard for diagnosis of hepatotoxicity,[67] non-invasive techniques such as magnetic resonance imaging/angiography (MRI/MRA) and Doppler ultrasound examination are frequently used as screening tools. A study comparing MRI to liver biopsy results found a sensitivity of 77% and a specificity of 72% in detecting NRH.[71] Furthermore, MRI is useful for detection of NRH-associated complications, including splenomegaly or ascites.[71] The European 6TG working party suggests screening by complete blood count, alanine aminotransferase, aspartate aminotransferase, alkaline phosphatase, gamma-glutamyl transferase, bilirubin, and C-reactive protein at baseline as well as one, two, four, eight, and 12 weeks followed by 3-month intervals.[70] When using low-dose 6TG, a liver biopsy is no longer routine practice but is reserved for patients with symptoms of portal hypertension or persisting liver test abnormalities.[43]

Determination of hepatic venous pressure gradient (HVPG) may be useful in the evaluation of the risk of complications, since varices only develop, when HVPG exceeds 10 mmHg. However, being an invasive technique it is rarely used.[31] The features of hepatotoxicity that have the highest sensitivity include discordant thrombocytopenia and splenomegaly. The latter is easily detected by clinical and ultrasound examinations. [72]

NRH is asymptomatic in most cases; leading to laboratory abnormalities without clinical signs, or is complicated by portal hypertension.[73] In a study evaluating 1,886 liver transplant patients, NRH was the reason for transplant in less than 1% of the cases.[74] In the studies included in this review, two patients required liver transplantation.[2,3] This indicates that NRH rarely leads to hepatic failure.

The present review emphasises the need for international standardised definitions of 6TG-related hepatotoxicity, clinical trials evaluating the use of non-invasive diagnostic methods in regard to detection of hepatotoxicity, and studies investigating the underlying mechanisms of 6TG-related hepatotoxicity. The included RCTs solely evaluated high doses of 6TG; hence, RCTs on low-dose 6TG are warranted. A RCT evaluating very low and individually titrated doses of 6TG (2.5–12.5 mg/m2/day) with concurrent conventional 6MP/methotrexate maintenance therapy in childhood ALL is currently being undertaken (ClinicalTrials.gov identifier:NCT02912676).

## Limitations

We included only patient populations with IBD and childhood ALL, which are the two largest groups of patients treated with 6TG. Generalisation in respect to 6TG-related hepatotoxicity of these two distinct populations must be interpreted with caution. A few separate reports exist of 6TG treatment in coeliac disease, psoriasis, autoimmune hepatitis, pilocytic astrocytoma, vertebral chloroma, and collagenous sprue. Furthermore, 6TG is used in the treatment of acute myeloid leukaemia as well as in intensive combination chemotherapy phases of many ALL protocols. These studies were not included because of short duration of 6TG administration and the extensive concomitant therapy with hepatotoxic drugs. Moreover, we found inconsistencies in the definitions of hepatotoxicity as well as in the diagnostic modalities across the included studies, which may limit the comparability between the studies. Finally, all studies investigating 6TG treatment in the IBD population were observational, without control groups, which entail higher risk of bias, particularly publication bias and confounding bias. However, observational studies are valuable for the description of rare adverse events, and provide a large body of evidence in the present review.

## Conclusions

Acute and severe hepatotoxicity occur in up to 25% of patients, when using 6TG doses of 40 mg/m^2^/day or higher in studies on 6TG for adult IBD and childhood ALL. 6TG-related hepatotoxicity persists in the form of NRH in 3% of the patients. However, the use of 6TG doses of approximately 12 mg/m^2^/day or less leads to hepatotoxicity in only 6% of the adult patients, corresponding to the incidence in the background population. Chronic hepatotoxicity in the form of portal hypertension has not been observed when using 6TG doses less than 40 mg/m^2^/day. Thus, 6TG doses below 12 /m^2^/day can be considered safe.

## Contributors

LNT and CUR are the guarantors of the review. LNT and KS conceptualised the review. LNT, RHL, TLF, KS and CUR drafted the protocol. LNT and CUR conducted the literature screening. LNT created and pilot tested the data extraction form. SA, MSS and LNT extracted the data, made assessments of risk of bias and confidence in cumulative evidence. LNT drafted the manuscript. The manuscript was read, revised, and approved by all authors.

## Acknowledgements

None.

## Supporting information

**S1 Appendix. Systematic review protocol S2 Appendix. Full search strategy**

**S3 Appendix. Detailed description of included studies**

**S4 Appendix. Risk of bias of included randomised controlled trials**

**S5 PRISMA 2009 Checklist**

